# The Gene Knockout of Angiotensin II Type 1a Receptor Improves High-fat Diet-Induced Obesity in rat *via* Promoting Adipose Lipolysis

**DOI:** 10.1101/2022.04.07.487533

**Authors:** Aiyun Li, Wenjuan Shi, Jin Wang, Xuejiao Wang, Yan Zhang, Zhandong Lei, Xiang-Ying Jiao

**Affiliations:** Key Laboratory of Cellular Physiology (Shanxi Medical University), Ministry of Education, and the Department of Physiology, Shanxi Medical University, Taiyuan 030001, China

**Keywords:** Angiotensin II type 1a receptor, obesity, adipose lipolysis, cAMP/PKA

## Abstract

**Aims:** The renin-angiotensin system (RAS) is over-activated and the serum angiotensin II (Ang II) level increased in obese patients, while their correlations were incompletely understood. This study aims to explore the role of Ang II in diet-induced obesity by focusing on adipose lipolysis.

**Methods:** Rat model of AT1aR gene knockout were established to investigate the special role of Ang II. Wild-type (WT) and AT1aR gene knockout (AT1aR^-/-^) SD rats were fed with normal diet or high-fat diet for 12 weeks. Adipose morphology and adipose lipolysis were examined.

**Results:** AT1aR deficiency activated lipolysis-related enzymes and increased the levels of NEFAs and glycerol released from adipose tissue in high-fat diet rats, while did not affect triglycerides synthesis. Besides, AT1aR knockout promoted energy expenditure and fatty acids oxidation in adipose tissue. cAMP levels and PKA phosphorylation in the adipose tissue were significantly increased in AT1aR^-/-^ rats fed with high-fat. Activated PKA could promote adipose lipolysis and thus improved adipose histomorphology and insulin sensitivity in high-fat diet rats.

**Conclusions:** AT1aR deficiency alleviated adipocyte hypertrophy in high-fat diet rats by promoting adipose lipolysis via cAMP/PKA pathway, and thereby delayed the onset of obesity and related metabolic diseases.

## 1. Introduction

Obesity is a growing health problem that induces major metabolic disorders, such as diabetes, cardiovascular disease, and hypertension. Obesity has been officially listed as a disease by the World Health Organization in 2000. It has been predicted that more than 57.8% adults and about 3.3 billion people worldwide will facing overweight or obesity problems by 2030[1]. With an increasing prevalence of obesity, it is urgent to elucidate the pathophysiological mechanism for obesity and develop effective therapy strategies for treatment of obesity and its associated disorders.

Adipose tissue plays major role in the development of obesity. Excess energy is stored in the adipose tissue in the form of triacylglycerol (TAG), while fatty acid mobilization via lipolysis releases fatty acids to increase hepatic fatty acid oxidation for energy supply[2]. Physiologically, adipose tissue maintains a dynamic balance between lipid synthesis and decomposition. However, adipose tissue stores excess energy by expansion and remodeling when the storage capacity of adipocytes is exceeded in response to overfeeding, resulting in fat accumulation and adipocyte hypertrophy[3, 4]. Therefore, inhibiting adipocyte hypertrophy can be an effective strategy for obesity treatment.

The renin-angiotensin system (RAS) is a dynamic physiologic system and its classic function is to regulate blood pressure and fluid and electrolyte balance. Angiotensin-converting enzyme inhibitor (ACEI) and angiotensin II receptor blockers (ARB) has been clinically used as antihypertensive drugs and play a crucial role in the treatment of cardiovascular diseases such as hypertension and heart failure. However, it has been found that these drugs have a positive effect in metabolic diseases such as obesity, insulin resistance and diabetes^[5, 6]^, indicating the relations between RAS and metabolic diseases. Animal studies have shown that mice with renin gene knockout are resistant to diet-induced obesity[7]. Besides, excessive activation of RAS is a common feature in obese patients[8, 9], and the serum levels of angiotensinogen (AGT) and angiotensin II (Ang II) in the obese are higher than those in normal population^[10-12]^. Adipose tissue is the most abundant source of AGT outside liver. It is showed that AGT expression increased in the adipose tissue of obese animal models and adipocyte-specific enhancement of AGT lead to insulin resistance[13], indicating the special role of adipose tissue RAS in regulation of metabolic homeostasis. Adipose RAS has attracted more and more attention due to its close relationship with obesity and adipose dysfunction over the years[14-16]. However, the underline mechanism by which RAS activation in adipose tissue results in obesity and other metabolic diseases remain unclear.

Angiotensin II (Ang II), the predominant peptide of the RAS, exerts effects mainly by binding with angiotensin type 1 receptor (AT1R) in adipose tissue[14]. Hence, we established an obese rat model with AT1aR gene knockout to explore the role of adipose RAS in the develop process of obesity. The results showed that the gene knockout of AT1aR ameliorated adipocyte hypertrophy by promoting adipose lipolysis, and thereby improving obesity and related metabolic disorders. Our finding suggests that targeting RAS signaling in adipose tissue may become a promising therapy avenue for obesity and associated metabolic abnormalities.

## 2. Materials and Methods

### 2.1 Experimental animals

Wild type male Sprague-Dawley (SD) rats purchased from the Experimental Animal Center of Shanxi Medical University and AT1aR^-/-^ male SD rats (Nanjing University-Nanjing Institute of Biology) were all fed in the SPF laboratory animal environmental facilities with 12 h light/dark cycles under standard room temperature (22 ± 2°C) and free access to water and food. AT1aR gene knockout rats have been verified as homozygous by polymerase chain reaction (PCR) (S1 Fig). All animal experiments were in accordance with the guidelines for the management of animals for medical experiments issued by the Ministry of Health of the People’s Republic of China (No. 55) and animal ethics standards of Shanxi Medical University, and approved by the ethics committee.

4-week male wild type (WT) rats and AT1aR^-/-^ rats were randomly divided into normal diet group (ND) and high-fat-diet group (HFD). The HFD group rats were fed with 60% high-fat feed (D12492, Whitby Technology Co., Ltd. Beijing,China) and the ND group rats were fed with normal feed for 12 weeks. Body weight was recorded weekly. At the end of feeding, rats were fasted overnight and anesthesia, blood sample was collected from the abdominal aorta for the assay of blood glucose, nonesterified free fatty acids (NEFAs), triglyceride (TG), total cholesterol (T-CHO), low-density lipoprotein (LDL) and high-density lipoprotein (HDL) contents with commercial kits (Jiancheng Bioengineering Institute, Nanjing, China). Serum glycerol content was measured with glycerol (liquid sample) enzymatic determination kit (Applygen Gene Technology Co., Ltd, Beijing). Epididymal adipose tissue were isolated immediately or stored at minus 80 degrees Celsius until analysis. Epididymal fat index were calculated as the ratio of epididymal fat weight to the body weight.

### 2.2 Glucose tolerance test and insulin tolerance test

For glucose tolerance test, rats were fasted overnight and administrated with glucose (2 g/kg) by gavage. For insulin tolerance test, rats were fasted for 6h and intraperitoneally injected insulin (1 IU/kg). Blood glucose contents at 0 min, 15 min, 30 min, 60 min, 90 min, and 120 min were recorded and blood glucose area under curve (AUC) was calculated.

### 2.3 Blood pressure measurement

At the end of 12-week feeding, rats were anesthetized and blood pressure were measured by carotid artery cannulation and monitored with the BL-420 system.

### 2.4 Morphological examination of adipose tissue

Isolated epididymal adipose tissue was fixed with 4% paraformaldehyde, dehydrated with gradient alcohol and then embedded in paraffin. Embedded wax block was sliced and epididymal sections were stained with hematoxylin for 20 min and with eosin for 15 min, respectively. After sealing slide with neutral gum, the morphology changes of adipocytes were visualized using optical microscope (Olympus, Japan) and the cross-sectional area of adipocytes were calculated with Image J software.

### 2.5 Quantitative real time RT-PCR

Total mRNA of epididymal adipose tissue was extracted with Trizol (Takara Bio Inc., Japan). The concentration and purity of extracted total mRNA were measured using NANODROP ONE (Thermo Scientific, USA). cDNA was synthesized with the PrimeScript™RT reagent kit (Takara Bio Inc., Japan) following the manufacturers’ instructions. Relative quantitative PCR was conducted with a TB Green Primer Ex Taq II (Takara Bio Inc., Japan) using LightCycler® 96 Real-Time PCR System (Roche, USA). The primer sequences were obtained from Takara and listed in supplementary Table 1. Gene expressions were normalized with β-actin. Statistically relative quantification was analyzed with equation 2^-ΔΔCT^. Ct is the threshold cycle to detect fluorescence.

**Table 1.**
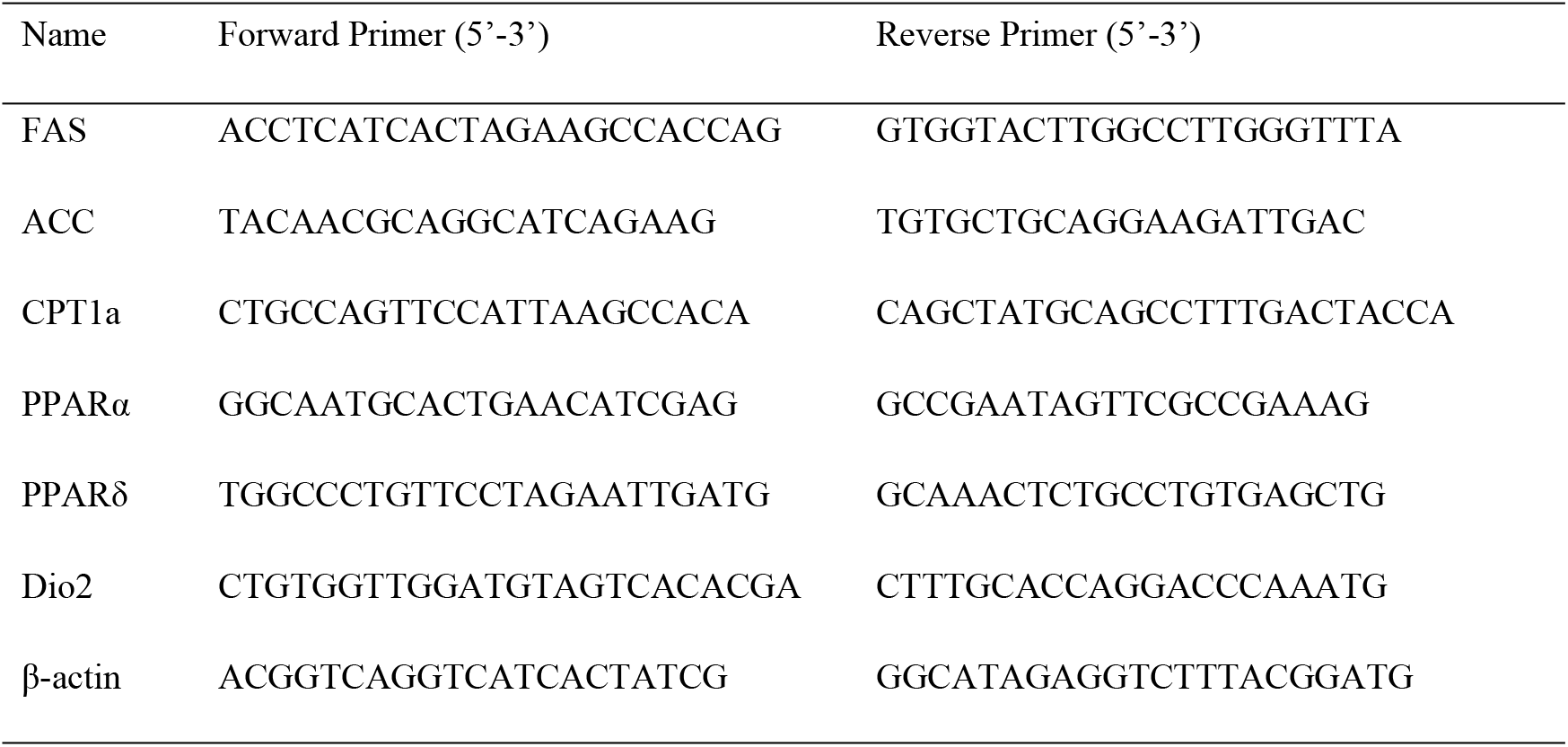
Primer sequences used for quantitative real time RT-PCR

### 2.6 Measurement of free fatty acids and glycerol in adipose tissue

At the end of 12-week feeding, rats were anesthetized and sterilized with alcohol. Epididymal adipose tissue was isolated and 200 mg adipose tissue was weight and cut into pieces, and then incubated in serum-free DMEM for 24 hours. DMEM was collected to measure NEFAs and glycerol contents in the supernatant with commercial Kits following the manufacturer’s instructions.

### 2.7 Measurement of cAMP concentration

Epididymal adipose tissue supernatants were obtained by homogenate and centrifugation. Cyclic AMP in the supernatants were measured using cAMP assay kit (4339, Cell Signaling Technology) according to the manufacturer’s protocol.

### 2.8 Western blot analysis

Epididymal adipose tissue was homogenized with a tissue grinder at 4°C temperature and the lysate were centrifugation at 12000 rpm for 20 minutes. The protein concentrations were measured with BCA kit (KeyGEN BioTECH Corp., Ltd, Nanjin, China). The protein samples were denatured, separated by SDS-PAGE and then transferred to the PVDF membrane. Membranes were blocked with 5% skim milk or bovine serum albumin (BSA) for 3 hours, and then incubated overnight at 4°C with specific primary antibodies as follows: ATGL (2138S, Cell Signaling Technology), P-HSL ^(ser660)^ (4126S, Cell Signaling Technology), HSL (4107S, Cell Signaling Technology), P-GSK-3β ^(ser9)^ (5558S, Cell Signaling Technology), GSK-3β (9315S, Cell Signaling Technology), β-actin (AP0060, Bioworld) and PKA (5842S, Cell Signaling Technology). Membranes were washed and incubated with secondary antibodies (BA1054, BOSTER Biological Technology) at room temperature for 3 hours. After washing, membranes were exposed by Super ECL Prime (SEVEN BIOTECH) with ChemiDoc™ Imaging System (BIO-RAD, USA). The images were analyzed quantitatively by densitometry with Image J software.

### 2.9. Statistical analysis

Results were shown as mean ± SEM. Data were analyzed by using Student’s t test for comparison the difference of two groups and the comparison among multiple groups was analyzed by one-way ANOVA. GraphPad Prism 6 was used for statistical analysis. A value of *p* < 0.05 was considered statistically significant.

## 3. Results

### 3.1 AT1aR knockout improved insulin sensitivity and metabolic disorders in high-fat diet rats

Glucose tolerance test and insulin tolerance test are important indicators that generally to be used to measure glucose tolerance and insulin sensitivity, respectively[17]. In response to glucose load or insulin injection, there was no significant differences between AT1aR^-/-^ rats and WT rats with normal diet, while the area under the curve (AUC) in HFD-fed AT1aR^-/-^ rats was significantly lower than that in the HFD-fed WT rats (Fig 1A, B), indicating that AT1aR knockout improved insulin sensitivity in high-fat-diet rats. Consistent with this, AT1aR knockout reduced the fasting blood glucose compared with WT rats fed with high-fat-diet (Fig 1C). Moreover, rats blood pressure was monitored through carotid artery intubation and the result showed that AT1aR gene knockout could reduce hypertension caused by high-fat feeding (Fig 1D).

**Fig 1.**
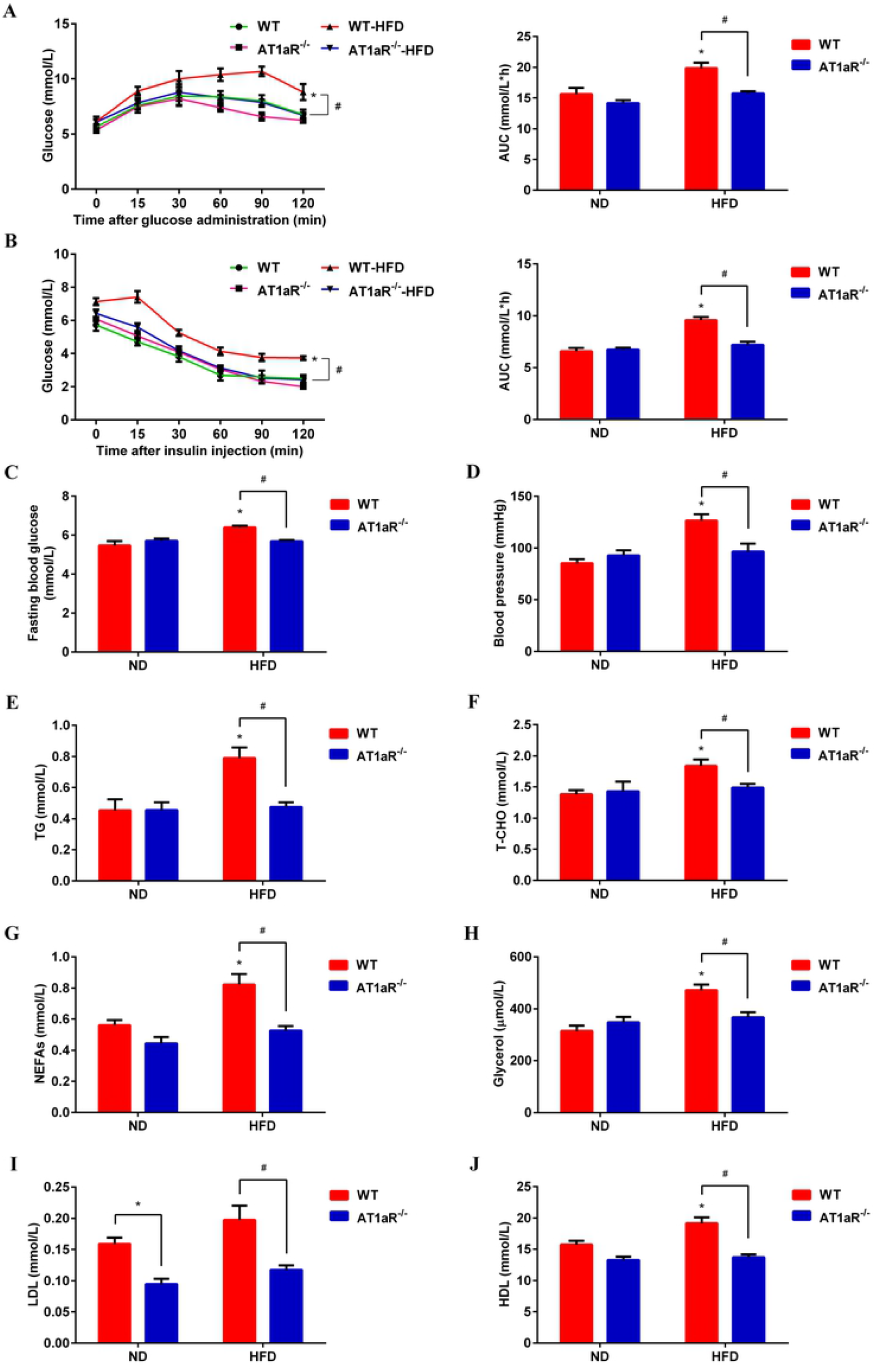
AT1aR knockout improved insulin resistance and metabolic disorders in high-fat diet rats. A: Oral glucose tolerance test (OGTT) curve and area under the curve. B: Insulin tolerance test (ITT) curve and area under the curve. C: Fasting blood glucose (FBG). D: Blood pressure. E: Serum triglyceride (TG) levels. F: Serum total cholesterol (T-CHO). G: Serum free fatty acid. H: Serum glycerol. I: Serum low-density lipoprotein (LDL). J: Serum high-density lipoprotein (HDL). *p < 0.05 vs WT-ND rats; #p < 0.05 vs WT-HFD rats. Data were presented as Mean ± S.E.M. n = 6

Next, lipid metabolism-related indicators in serum were detected with commercial kits. The results showed that serum levels of TG, T-CHO, NEFAs, glycerol, LDL and HDL in high-fat diet WT rats were significantly higher than those of WT rats fed with normal diet, whereas these alterations were significantly reversed by AT1aR gene knockout, demonstrating that AT1aR knockout improved metabolic disorders in obese rats (Fig 1E-J). Besides, serum LDL level of AT1aR^-/-^ rats were also lower than that of the WT rats fed with normal diet (Fig 1I).

### 3.2 AT1aR knockout alleviated high-fat diet-induced adipocyte hypertrophy

Body weight of rats were recorded weekly and the results showed that body weight gain of WT rats were increased by high-fat diet feeding, while body weight gain of high-fat diet AT1aR^-/-^ rats were much lower than that of high-fat diet WT rats (Fig 2A, B). Epididymal adipose tissues were isolated and their physical morphology were photographed. As showed in Fig 2C, physical morphology of epididymal adipose tissue in AT1aR^-/-^ rats was smaller than that of WT rats. Besides, the epididymal fat index of high-fat diet AT1aR^-/-^ rats was also decreased compared to high-fat diet WT rats (Fig 2D). Moreover, the histomorphology of adipose were visualized by HE staining and adipocyte area of WT rats were significantly enlarged by high-fat diet feeding, whereas this alteration were significantly reversed in high-fat diet AT1aR^-/-^ rats (Fig 2E). These results indicated that AT1aR gene knockout improved adipose histomorphology and inhibited adipocyte hypertrophy.

**Fig 2.**
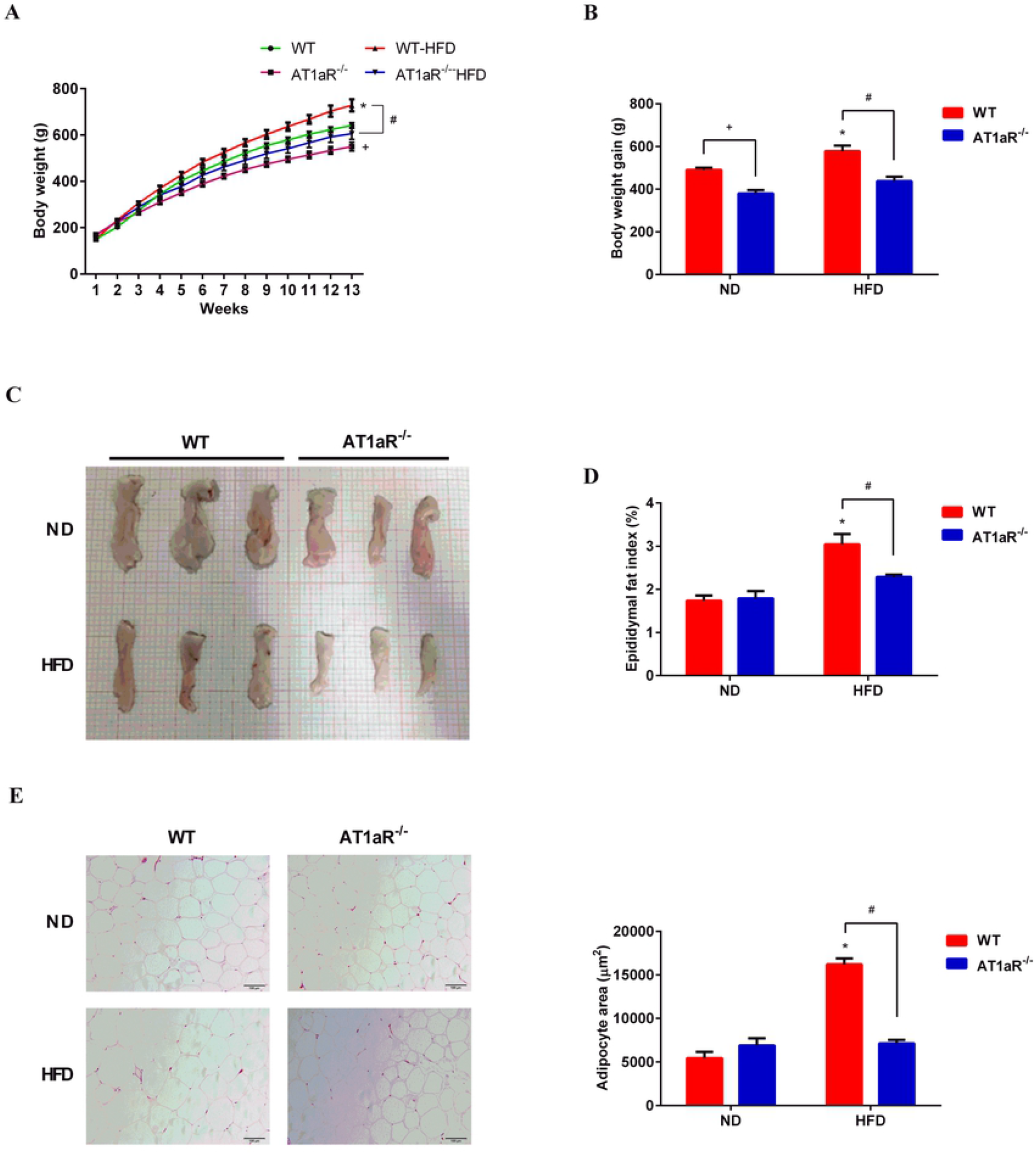
AT1aR knockout improved epididymal histomorphology induced by high fat diet. A, B: Body weight of rats was recorded weekly and body weight gain was calculated by weight increase in 12 weeks. C: Physical morphology of epididymal fat; D: Epididymal fat index (showed as epididymal fat weight/body weight). E: Adipocyte morphology was visualized by HE staining and adipocyte area were calculate by Image J. *p < 0.05 vs WT-ND rats; #p < 0.05 vs WT-HFD rats. Data were presented as Mean ± S.E.M. n = 6

### 3.3 AT1aR gene knockout promoted adipose lipolysis

The main function of white adipose tissue is to store energy in the form of triglycerides and the fat mass is systematically regulated by lipid synthesis and decomposition. In order to know the underlining mechanism by which AT1aR gene knockout inhibited adipocyte hypertrophy, we detected the expressions of major enzymes for triglyceride synthesis and lipolysis in the adipose tissue. Fatty acid synthase (FAS) and acetyl-CoA carboxylase (ACC) are rate-limiting enzymes involved in biosynthesis of fatty acids, an important step of lipogenesis[18]. Gene expressions of FAS and ACC were measured with QPCR and the results showed that there were no significant differences between WT and AT1aR^-/-^ rats neither fed with normal diet nor with high-fat diet (Fig 3A, B). The hydrolysis of triglycerides is initiated by adipose triglyceride lipase (ATGL)[19], and hormone sensitive lipase (HSL) is the main hydrolase for triacylglycerol (TAG) and diacylglycerol (DAG)[20]. As showed in Fig 3C, protein expression of ATGL and phosphorylation of HSL in AT1aR^-/-^ rats were much higher than those of WT rats both in the normal and high-fat diet rats. Furthermore, the adipose tissue was cultured in the DMEM for 24 hours and the levels of NEFAs and glycerol in the culture medium were detected. Compared with WT rats fed with high-fat diet, the levels of NEFAs and glycerol released from adipose tissue of high-fat diet AT1aR^-/-^ rats increased significantly (Fig 3D, E). These results proved that AT1aR knockout improved adipose histomorphology in high-fat diet rats by promoting adipose lipolysis, while did not affect lipid synthesis.

**Fig 3.**
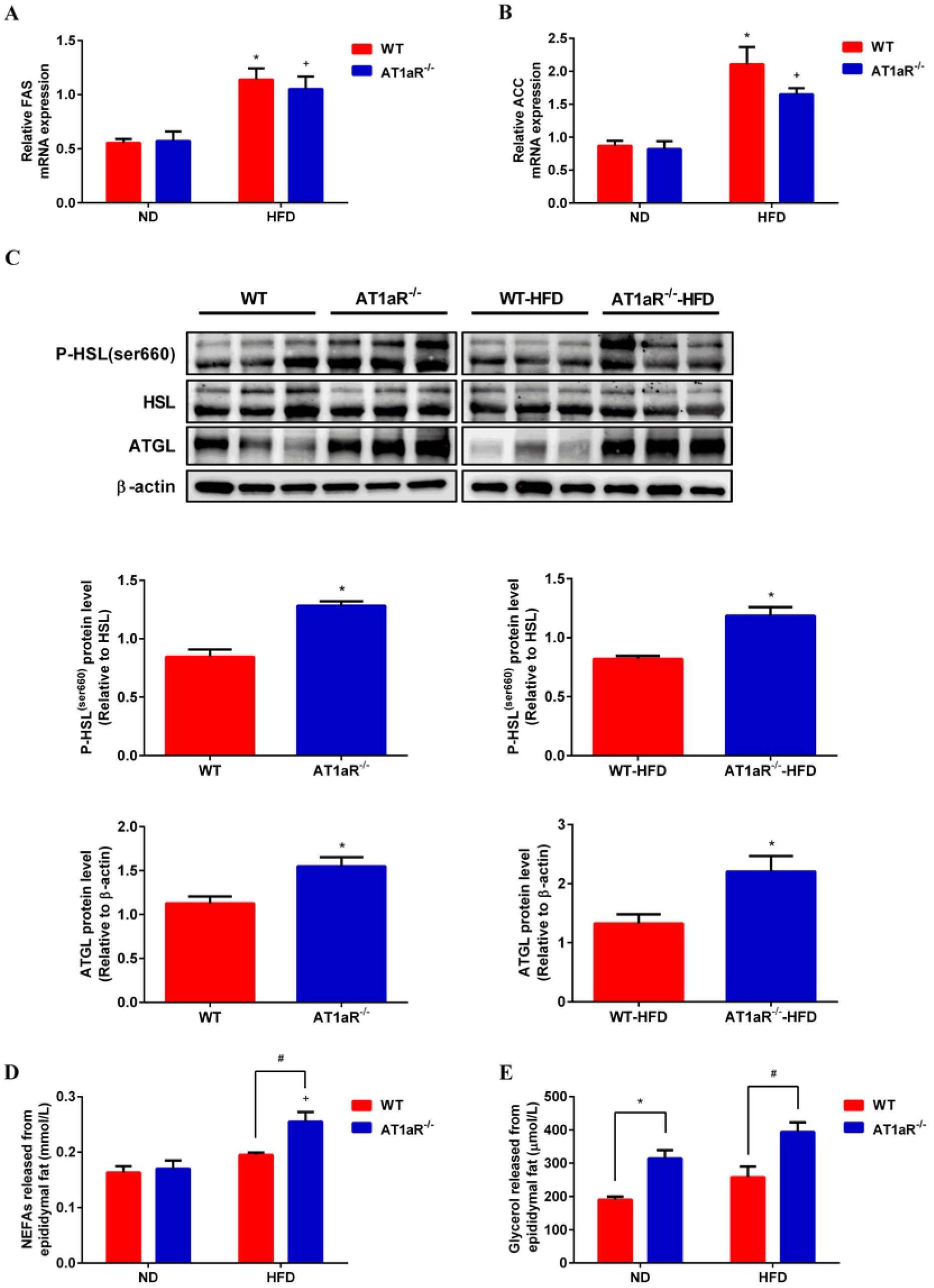
AT1aR knockout promoted adipose lipolysis. A, B: Gene expressions of key enzymes for lipid synthesis in adipose tissue, FAS (A) and ACC (B). C: Protein expressions of adipose lipolysis related enzymes, ATGL and HSL. D, E: Levels of NEFAs (D) and glycerol (E) in culture medium released by adipose tissue within 24h. *p < 0.05 vs WT-ND rats; +p < 0.05 vs AT1aR-/--ND rats. Data were presented as Mean ± S.E.M. n = 6

### 3.4 AT1aR knockout accelerated adipose energy expenditure and fatty acids oxidation

Subsequently, adipose lipid utilization was detected. Peroxisome proliferator activated receptor δ (PPARδ) is a transcription factor that promote oxidative metabolism in adipose[21] and deiodinase 2 (Dio2) could activate thyroid hormone to promote energy expenditure[22]. The results showed that by comparison of WT rats fed with high-fat diet, the gene expressions of PPARδ and Dio2 in adipose tissue were significantly increased in high-fat diet AT1aR^-/-^ rats (Fig 4A, B), indicating increased energy consumption adipose tissue of AT1aR^-/-^ rats. Besides, the levels of fatty acid oxidation in adipose tissue were also measured. Peroxisome proliferator-activated receptors α (PPARα) acts as a transcription factor to regulate a series of genes involved in fatty acid oxidation like carnitine palmitoyltransferase 1 (CPT1) [23, 24]. The results showed that the gene expressions of PPARα and CPT1 were significantly increased in AT1aR^-/-^ rats compared with WT rats fed with high fat diet. However, there was not difference between the WT and AT1aR^-/-^ rats fed a normal diet (Fig 4C, D). These results indicated enhanced fatty acid oxidation in high-fat diet AT1aR^-/-^ rats.

**Fig4.**
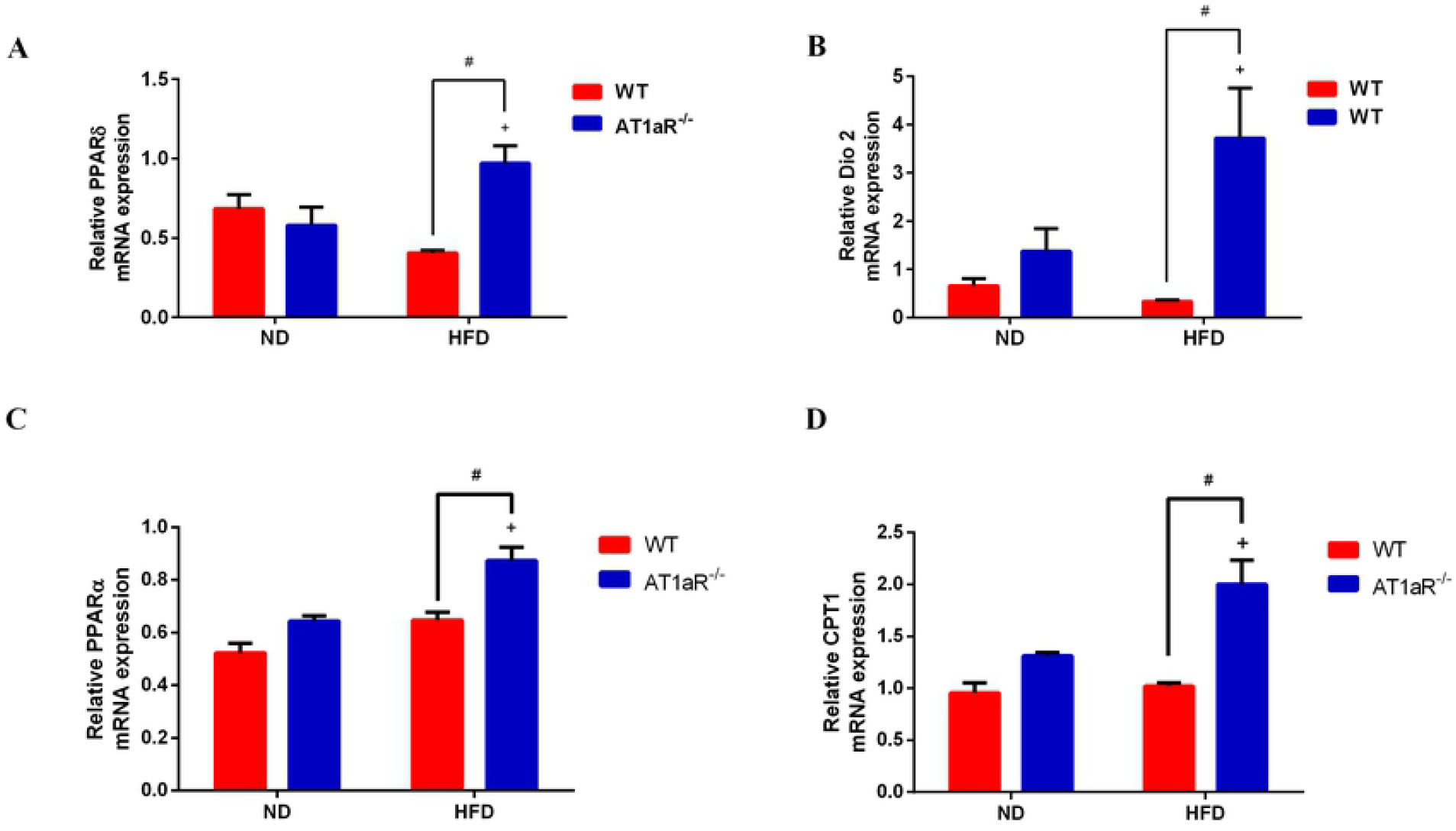
AT1aR knockout accelerated adipose energy expenditure and fatty acids oxidation. A, B: Gene expressions of PPARδ (A) and Dio2 (B) in adipose tissue. C, D: Gene expressions of PPARα and CPT1 in adipose tissue. +p < 0.05 vs AT1aR-/--ND rats. Data were presented as Mean ± S.E.M. n = 6

### 3.5 AT1aR knockout activated cAMP/PKA pathway

It has been reported that the binding of Ang Ⅱ to AT1R activates Gi protein against the effect of cAMP/PKA pathway[25]. As showed in Fig 5A and 5B, PKA and cAMP levels were significantly increased in AT1aR^-/-^ rats fed with high-fat. Protein kinase A (PKA) that activated by cyclic adenosine monophosphate (cAMP) mainly phosphorylates HSL to mediate adipose lipolysis [26]. Apart from HSL, GSK-3β is also an important phosphorylation substrate of PKA[27]. The phosphorylation level of GSK-3β in AT1aR^-/-^ rats was higher than WT rats both in the normal and high-fat diet (Fig 5A), further confirmed the enhancement of PKA activity in AT1aR^-/-^ rats. These results suggested that gene knockout of AT1aR could activate cAMP/PKA pathway.

**Fig 5.**
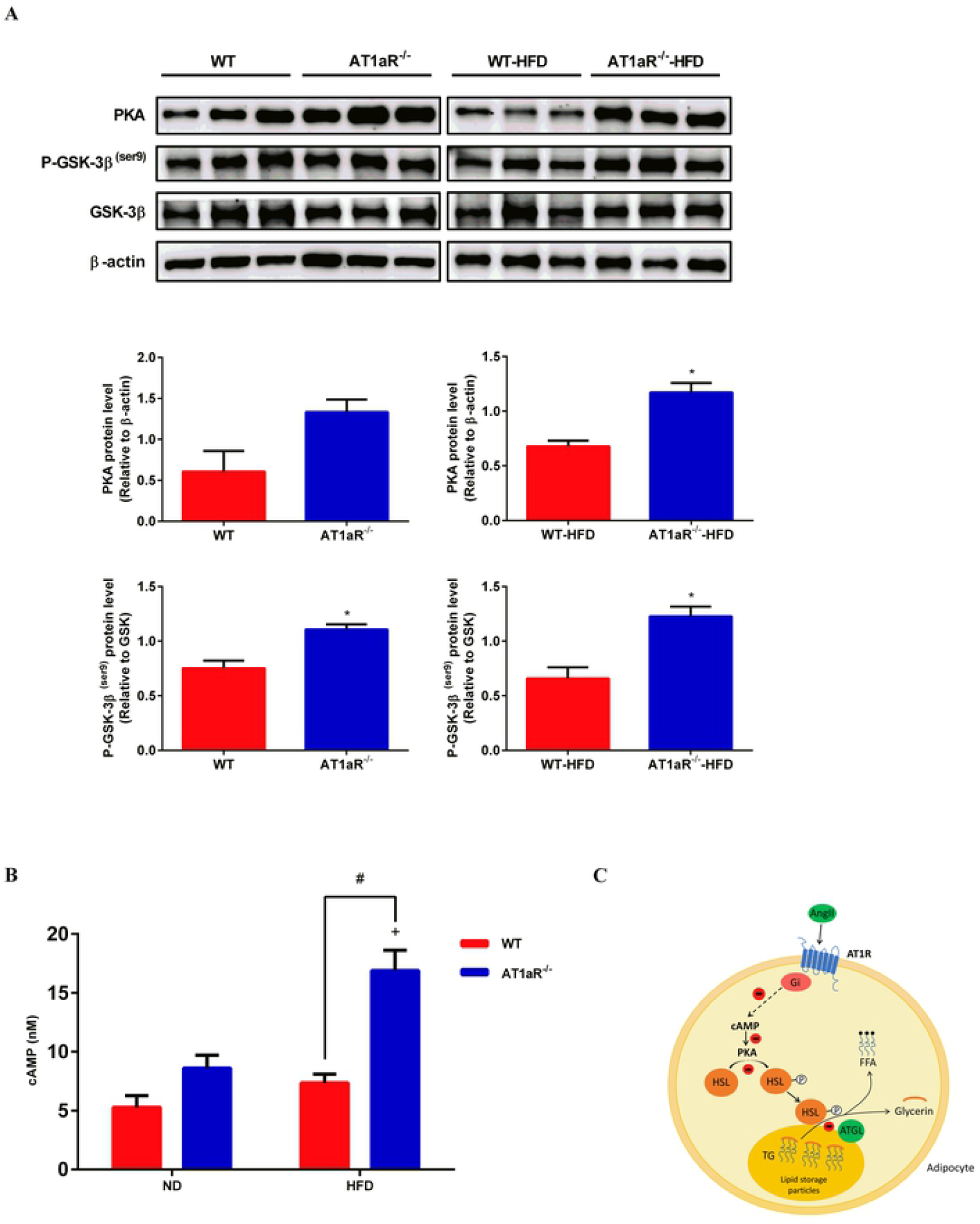
AT1aR knockout activated cAMP/PKA pathway. A: Protein expressions of PKA, P-GSK-3β (ser9) and GSK-3β in adipose tissue. B: cAMP levels in adipose tissue. C: The proposed pathway for Ang II inhibiting adipose lipolysis. By binding to AT1R, Ang II activates inhibitory Gi which in turn reduces cAMP production. As a result, PKA activation is limited and HSL phosphorylation is decreased, leading to inhibitory triglyceride hydrolysis. +p < 0.05 vs AT1aR-/--ND rats. Data were presented as Mean ± S.E.M. n = 6

## 4. Discussion

Obesity is the common pathological basis of many metabolic diseases and its mechanism exploration has attracted more and more attention. RAS is generally considered acting in the regulation of blood pressure and organism water-salt balance. However, clinical studies have demonstrated that type 2 diabetes can be delayed by treatment with ACEI and ARB compared with other antihypertensive drugs[28]. Besides, captopril, one of ACEI drugs, was showed to not only lower blood pressure but also reduce the weight of high-fat diet rats[29]. These studies indicated that RAS plays an important role in obesity and subsequent metabolic diseases, whereas the regulatory mechanism remains unclear. In the present study, we found that the gene knockout of AT1aR improved high-fat diet induced obesity by promoting lipolysis through cAMP/PKA pathway, providing new sight for the prevention and treatment of obesity and related metabolic disorders.

As the primary peptide of the RAS, Ang II exerts its biological activity mainly through AT1R in adipose tissue. Different from human beings, AT1R is coded by two gene subtypes, AT1aR and AT1bR, in rodent mammals[30]. AT1aR has the most homology with human, mainly involved in the vasoconstriction and blood pressure regulation. While AT1bR is related to the thirst response of mammals and existed in certain areas of the central nervous system and adrenal tissues [31]. So, in the present study, we constructed AT1aR^-/-^ rats to explore the relation between RAS and obesity.

Obesity is closely correlated with insulin resistance and diabetes. Besides storage energy, adipose tissue also acts as an endocrine organ. In obesity, adipocytes secrete inflammatory adipokines which damage insulin signaling pathway, resulting in disorders of glucose and lipid metabolism^[32]^. Here, we showed that AT1aR gene knockout improved glucose intolerance and insulin sensibility in high-fat diet induced obese rats. Consistent with our results, Kengo Azushima et al. has demonstrated that upregulation of AT1R-related protein (ATRAP), a protein limiting AT1R’s effects by promoting its internalization, could improve insulin resistance[17].

Lipid stocks in the adipose tissue fluctuate by lipid synthesis and lipolysis. Our results showed that AT1aR gene knockout improved adipocyte hypertrophy in obese rats mainly by promoting lipolysis. However, enhanced lipolysis increases circulatory NEFAs which leading to ectopic deposition of lipids. Interestingly, serum levels of NEFAs were decreased in high-fat diet AT1aR^-/-^ rats despite significant enhancement of lipolysis in adipose tissue. In this regard, energy expenditure and NEFAs oxidation in adipose tissue were measured. The results demonstrated that genes expressions of proteins involved in adipose tissue oxidative metabolism were elevated. Given that increasing lipolysis in adipose tissue result in elevated circulating NEFAs, our results showed that AT1aR gene knockout promoted adipose energy expenditure and fatty acids utilization, and thus lowering lipolysis-derived circulating NEFAs levels.

By binding to AT1R, a G protein-coupled receptor, Ang Ⅱ activates inhibitory Gi protein which in turn decreases cAMP production[33, 34]. The cAMP/PKA pathway plays an important role in energy balance and metabolic regulation. As an important second messenger, increased cellular cAMP activates PKA which in turn phosphorylates HSL[35]. AT1aR gene knockout increased PKA protein expression and cellular levels of cAMP in the adipose tissue. Activated PKA can be confirmed by increased phosphorylation of GSK-3β, a main phosphorylation substrate of PKA[27].

In conclusion, our results showed that RAS predominant peptide Ang II promoted adipocyte hypertrophy by inhibition of cAMP/PKA and subsequent adipose lipolysis via binding to AT1aR. However, AT1aR gene knockout activated cAMP/PKA signaling and elevated adipose lipolysis by phosphorylation of HSL, and thus improving obesity and insulin resistance (Fig 5C). Our findings emphasized the special role of RAS signaling in adipose tissue in the progress of obesity and associated metabolic abnormalities, and provided potential drug target for their therapy.

## Acknowledgments

Thanks for the support of Shanxi Key Subjects Construction (FSKSC), and Shanxi ‘1331 Project’ Key Subjects Construction.

## Author contributions

**Aiyun Li:** Conceptualization, Methodology, Writing - Review & Editing. **Wenjuan Shi:** Conceptualization, Methodology, Investigation, Writing - Original Draft. **Jin Wang:** Resources, Project administration, Validation. **Xuejiao Wang:** Investigation, Software. **Yan Zhang:** Validation, Software. **Zhandong Lei:** Data Curation, Software. **Xiangying Jiao:** Conceptualization, Writing - Review & Editing, Supervision.

## Supporting information

**S1 Fig.**
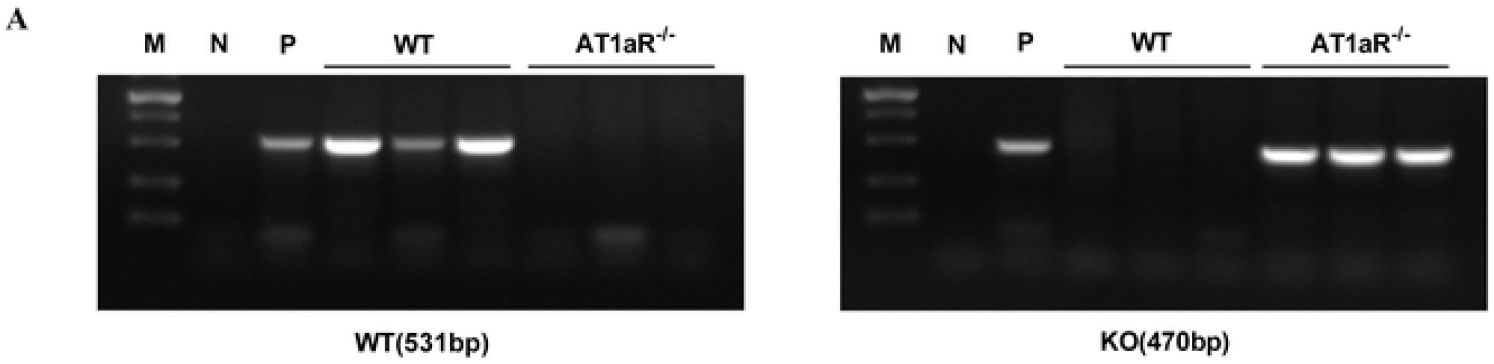
PCR results for AT1aR-/- rats and WT rats. M: marker; N: negative control; P: positive control

